# Genome sequencing of turmeric provides evolutionary insights into its medicinal properties

**DOI:** 10.1101/2020.09.07.286245

**Authors:** Abhisek Chakraborty, Shruti Mahajan, Shubham K. Jaiswal, Vineet K. Sharma

## Abstract

*Curcuma longa*, or turmeric, is traditionally known for its immense medicinal properties and has diverse therapeutic applications. However, the absence of a reference genome sequence is a limiting factor in understanding the genomic basis of the origin of its medicinal properties. In this study, we present the draft genome sequence of *Curcuma longa*, the first species sequenced from Zingiberaceae plant family, constructed using 10x Genomics linked reads. For comprehensive gene set prediction and for insights into its gene expression, the transcriptome sequencing of leaf tissue was also performed. The draft genome assembly had a size of 1.24 Gbp with ~74% repetitive sequences, and contained 56,036 coding gene sequences. The phylogenetic position of *Curcuma longa* was resolved through a comprehensive genome-wide phylogenetic analysis with 16 other plant species. Using 5,294 orthogroups, the comparative evolutionary analysis performed across 17 species including *Curcuma longa* revealed evolution in genes associated with secondary metabolism, plant phytohormones signaling, and various biotic and abiotic stress tolerance responses. These mechanisms are crucial for perennial and rhizomatous plants such as *Curcuma longa* for defense and environmental stress tolerance via production of secondary metabolites, which are associated with the wide range of medicinal properties in *Curcuma longa*.

## INTRODUCTION

Turmeric, a common name for *Curcuma longa*, is traditionally used as a herb and spice since 4,000 years in Southern Asia [1]. It has a long history of usage in medicinal applications, as an edible dye, as a preservative in many food materials, in religious ceremonies, and is now widely used in cosmetics throughout the world [1], [2]. It is a perennial rhizomatous monocot herb belonging to the *Curcuma* genus that contains ~70 species belonging to the family Zingiberaceae that comprises of more than 1,300 species, which are known to be widely distributed in tropical Africa, Asia and America [3], [4]. This family is enriched in rhizomatous and aromatic plants with a variety of bioactive compounds. Zingiberaceae family members are known to be associated with endophytes, which produce various secondary metabolites in the species of Zingiberaceae family, thus in turn confer medicinal properties [4].

Secondary metabolism is one of the key adaptations in plants to cope with the environmental conditions through the production of a wide range of common or plant-specific secondary metabolites [5], [6]. These secondary metabolites also play a key role in plant defense mechanisms, and several of these metabolites have numerous pharmacological applications in herbal medicines and phytotherapy [7]. The pathways for biosynthesis of various secondary metabolites including phenylpropanoids, flavonoids (such as curcuminoids), terpenoids, and alkaloids are found in *Curcuma longa* [8]–[12], and also possesses other constituents such as volatile oils, proteins, resins and sugars [13]. Flavonoids are known to have anti-inflammatory, antioxidant, and anti-cancer activities [14], [15]. Phenylpropanoids are also of great importance because of their antioxidant, anti-cancer, anti-microbial, anti-inflammatory, and wound-healing activities [16]. The other class of secondary metabolites, terpenoids, are also known to possess anti-cancer and anti-malarial properties [17].

The three curcuminoids namely curcumin, demethoxycurcumin, and bisdemethoxycurcumin, are responsible for the yellow colour of turmeric [13]. Among these, the primary bioactive component of turmeric is curcumin, which is a polyphenol-derived flavonoid compound and also known as diferuloylmethane [13]. Curcumin has been known to show broad-spectrum antimicrobial properties against bacteria, fungi and viruses [18], and also possesses anti-diabetic, anti-inflammatory, antifertility, anti-coagulant, hepatoprotective, and hypertension protective properties [13], [19]. Being an excellent scavenger of reactive oxygen species (ROS) and reactive nitrogen species [20], its antioxidant activity also controls DNA damage by lipid peroxidation mediated by free-radicals, and thus provides it with anti-carcinogenic properties [20], [21]. Due to these medicinal properties, turmeric has been of interest for scientists from many decades.

Several studies have been carried out to study the secondary metabolites and medicinal properties of this plant [22], [23]. Recently, the transcriptome profiling and analysis of *Curcuma longa* using rhizome samples have been carried out to identify the secondary metabolite pathways and associated transcripts [11], [17], [24], [25]. However, its reference genome sequence is not yet available, which is much needed to understand the genomic and molecular basis of the evolution of the unique characteristics of *Curcuma longa*. According to the Plant DNA c-value database, the genome is polyploid with 2n=63 chromosomes and has an estimated size of 1.33 Gbp [26].

Therefore, we performed the first draft genome sequencing and assembly of *Curcuma longa* using 10x Genomics linked reads generated on Illumina platform. The transcriptome of rhizome tissue for this plant has been known from previous studies but the transcriptome of leaf tissue is yet unknown [11], [17], [24], [25]. Thus, we carried out the first transcriptome sequencing of leaf tissue and along with the available RNA-Seq data of rhizome, performed a comprehensive transcriptome analysis that also helped in the gene set construction. We also constructed a genome-wide phylogeny of *Curcuma longa* with other available monocot genomes. The comparative analysis of *Curcuma longa* with other monocot genomes revealed adaptive evolution in genes associated with plant defense and secondary metabolism, and provided genomic insights into the medicinal properties of this species.

## METHODS

### Sample collection, library preparation and Sequencing

The plant sample was collected from an agricultural farm (23.2280252°N 77.2088987°E) located in Bhopal, Madhya Pradesh, India. The leaves were homogenized in liquid nitrogen for DNA extraction using Carlson lysis buffer. Species identification was performed by PCR amplification of a nuclear gene (Internal Transcribed Spacer ITS) and a chloroplast gene (Maturase K), followed by Sanger sequencing at the in-house facility. The library construction from the extracted DNA was done with the help of Chromium Controller instrument (10x Genomics) using Chromium™ Genome Library & Gel Bead Kit v2 by following the manufacturer’s instructions. RNA extraction was carried out for the powdered leaves using TriZol reagent (Invitrogen, USA). The transcriptomic library was prepared with TruSeq Stranded Total RNA Library Preparation kit by following the manufacturer’s protocol with Ribo-Zero Workflow (Illumina, Inc., USA). The quality of libraries was evaluated on Agilent 2200 TapeStation using High Sensitivity D1000 ScreenTape (Agilent, Santa Clara, CA) prior to sequencing. The prepared genomic and transcriptomic libraries were sequenced on Novaseq 6000 (Illumina, Inc., USA) generating 150 bp paired-end reads. The detailed DNA and RNA extraction steps and other methodologies are mentioned in **Supplementary Text S1**.

### Genomic data processing and assembly

The barcode sequences were trimmed from raw 10x Genomics linked reads using a set of python scripts (https://github.com/ucdavis-bioinformatics/proc10xG). The genome size of *Curcuma longa* was estimated using a k-mer count distribution method implemented in SGA-preqc **(Supplementary Text S1)** [27]. A total of 631.11 million raw 10x Genomics linked reads corresponding to ~71X coverage were used for generating a *de novo* assembly using Supernova assembler v2.1.1 with maxreads=all option and other defaults settings [28]. The haplotype-phased assembled genome was generated using Supernova mkoutput in ‘pseudohap’ style.

The 10x Genomics linked reads were run through Longranger basic v2.2.2 (https://support.10xgenomics.com/genome-exome/software/pipelines/latest/installation) for barcode processing and were used to detect and correct mis-assemblies in Supernova assembled genome using Tigmint v1.1.2 [29]. The first round of scaffolding was carried out using ARCS v1.1.1 (default parameters) to generate more contiguous assembly using 10x Genomics linked reads [30]. Further scaffolding was performed to improve the contiguity using AGOUTI v0.3.3 with the quality filtered paired-end RNA-Seq reads, which were also used in *de novo* transcriptome assembly [31]. Gap-closing was performed with barcode-processed linked reads using Sealer v2.1.5 with k-mer value from 30 to 120 with an interval of 10 bp using a Bloom filter-based approach [32]. Finally, the assembly quality was improved by Pilon v1.23 using barcode-processed linked reads to fix small indels, individual base errors, or local mis-assemblies that could be introduced by the previous scaffolding steps [33]. BUSCO v4.0.4 was used to assess the genome assembly completeness using embryophyta_odb10 database [34]. The other details about the genome assembly post-processing are mentioned in **Supplementary Text S2**.

### Transcriptome assembly

The RNA-Seq data from our study and from previously available transcriptome studies of *Curcuma longa* were used for *de novo* transcriptome assembly [11], [17], [24], [25], [35]. Trimmomatic v0.38 was used for adapter removal and quality filtration of raw Illumina sequence data **(Supplementary Text S1)** [36]. Finally, Trinity v2.9.1 was used with default parameters to perform *de novo* transcriptome assembly of quality-filtered paired-end and single-end reads [37]. Assembly statistics was calculated using a Perl script available in Trinity software package. The longest isoforms for all genes were extracted from the assembled transcripts. These sequences were clustered to identify the unigenes using CD-HIT-EST v4.8.1 (90% sequence identity, 8 bp seed size) [38].

### Genome annotation

The final polished assembly (length ≥500 bp) was used for genome annotation. RepeatModeler v2.0.1 was used to generate a *de novo* repeat library for this genome [39]. The resultant repeat sequences were further clustered using CD-HIT-EST v4.8.1 (90% sequence identity, seed size = 8 bp) [38]. This repeat library was used to soft-mask *Curcuma longa* genome using RepeatMasker v4.1.0 (http://www.repeatmasker.org) and was further used for gene set construction. Genome annotation was carried out using MAKER pipeline to predict the final gene models using *ab initio* gene prediction programs and evidence-based approaches [40]. The transcriptome assembly of *Curcuma longa*, and protein sequences of *Curcuma longa* along with its closest species *Musa acuminata* were used as empirical evidence in MAKER pipeline. AUGUSTUS v3.2.3, BLAST and Exonerate v2.2.0 were used in MAKER pipeline for *ab initio* gene prediction, evidence alignments and alignments polishing, respectively [41], [42]. Tandem Repeat Finder (TRF) v4.09 was used to detect the tandem repeats present in *Curcuma longa* genome [43]. Additionally, miRBase database and tRNAscan-SE v2.0.5 were used for homology-based identification of miRNAs and *de novo* prediction of tRNAs, respectively [44], [45]. The other details about genome annotation are mentioned in **Supplementary Text S3**.

### Construction of gene set

The coding genes were predicted by TransDecoder v5.5.0 from the unigenes identified in transcriptome assembly, using Uniref90 and Pfam databases for homology-based searching as ORF retention criteria (https://github.com/TransDecoder/TransDecoder) [46]–[49]. Also, the MAKER pipeline-based gene models were clustered using CD-HIT-EST v4.8.1 (95% sequence identity, 8 bp seed size) [38]. The TransDecoder pipeline derived genes (≥300 bp) were aligned against the MAKER derived gene set (≥300 bp) using BLASTN, and the genes that did not match with identity ≥50%, query coverage ≥50% and e-value 10^-9^ were added to the MAKER gene set to prepare the final gene set of *Curcuma longa* [50].

### Orthogroups identification

Representative species from all 15 monocot genus available in Ensembl plants release 47, and model organism *Arabidopsis thaliana* as an outgroup species were selected for orthogroups identification [51]. To construct the orthogroups, the protein sequences of *Curcuma longa* obtained from TransDecoder and proteome files for other 16 selected species i.e., *Aegilops tauschii*, *Ananas comosus, Brachypodium distachyon, Dioscorea rotundata, Eragrostis tef, Hordeum vulgare, Leersia perrieri, Musa acuminata, Oryza sativa, Panicum hallii fil2, Saccharum spontaneum, Setaria italica, Sorghum bicolor*, *Triticum aestivum*, *Zea mays*, and *Arabidopsis thaliana* obtained from Ensembl release 47, were used. The longest isoforms for all proteins were extracted for all selected species to construct the orthogroups using OrthoFinder v2.3.9 [52].

### Construction of orthologous gene set and phylogenetic tree

Only those orthogroups that contained genes from all 17 species were extracted from all the identified orthogroups. The fuzzy one-to-one orthogroups containing genes from all 17 species were identified from these orthogroups, and extracted using KinFin v1.0 [53]. For cases where the orthologous gene sets comprised of multiple genes for a species, the longest gene was extracted. The fuzzy one-to-one orthogroups were further aligned individually using MAFFT v7.467 for species phylogenetic tree construction [54]. BeforePhylo v0.9.0 (https://github.com/qiyunzhu/BeforePhylo) was used to trim the multiple sequence alignments to remove empty sites and to concatenate the multiple sequence alignments of all fuzzy one-to-one orthologous gene sets across 17 species. This concatenated alignment was used by the rapid hill climbing algorithm-based RAxML v8.2.12 for construction of maximum likelihood species phylogenetic tree with ‘PROTGAMMAGTR’ amino acid substitution model using 100 bootstrap values [55].

### Identification of genes with higher nucleotide divergence

Protein sequences of all the orthogroups across 17 species were aligned individually using MAFFT v7.467 [54], and individual maximum likelihood-based phylogenetic trees were built using these alignments by RAxML v8.2.12 (‘PROTGAMMAGTR’ amino acid substitution model, 100 bootstrap values) [55]. The ‘adephylo’ package in R was used to calculate root-to-tip branch length distance values for each species [56]. *Curcuma longa* genes that showed comparatively higher root-to-tip branch length with respect to the other selected species were extracted, and were considered as the genes with higher nucleotide divergence or higher rate of evolution.

### Identification of genes with unique substitution having functional impact

The unique substitutions in genes that have impact on protein function can identify species-specific amino acid substitutions and are considered as a site-specific evolutionary signature. However, the inclusion of phylogenetically distant species in this analysis may erroneously increase the number of uniquely substituted genes, therefore we restricted this analysis by only considering the monocot species (available on the Ensembl plant release 47) for reliable results. The amino acid positions that were identical across the other 16 species in the individual multiple sequence alignments of all orthogroups but different in *Curcuma longa* were considered as the uniquely substituted amino acid positions.

For identification of uniquely substituted sites, an in-house python script was used. Any gap and ten amino acid sites around any gap in the alignments were not considered in this analysis. The impact of these unique amino acid substitutions on protein function was identified using Sorting Intolerant From Tolerant (SIFT), by utilizing UniProt as the reference database [57], [58].

### Identification of positively selected genes

SATé v2.2.7 was used to generate individual protein alignments for all orthogroups across 17 species [59]. For its iterative alignment approach, SATé uses PRANK, MUSCLE and RAxML for sequence alignment, merging the alignment, and tree estimation, respectively [55], [60], [61]. The ambiguous sites present in the codons were filtered and protein alignment-based nucleotide alignments were performed for protein-coding nucleotide sequences of all orthogroups across 17 species using ‘tranalign’ program of EMBOSS v6.6.0 package [62]. ‘codeml’ from PAML package v4.9a that uses a branch-site model was used to identify positively selected genes using nucleotide alignments of all the orthologs in phylip format and the species phylogenetic tree generated in previous steps [63]. Log-likelihood values were used to perform likelihood ratio tests and chi-square analysis-based p-values were calculated. The genes that qualified against the null model (fixed omega) (FDR-corrected p-values <0.05) were identified as positively selected genes. All codon sites showing greater than 95% probability for foreground lineage based on Bayes Empirical Bayes (BEB) analysis were termed as positively selected sites.

### Genes with multiple signs of adaptive evolution (MSA)

Among the three signs of adaptive evolution– higher nucleotide divergence, unique substitution having functional impact and positive selection, the *Curcuma longa* genes that showed at least two of these signs were termed as genes with multiple signs of adaptive evolution or MSA genes [64].

### Functional annotation

KAAS genome annotation server v2.1 was used to assign KEGG Orthology (KO) identifiers and KEGG pathways to the genes [65]. eggNOG-mapper v2 was used for functional annotation of genes using precomputed orthologous groups from eggNOG clusters [66]. WebGeStalt web server was used for GO enrichment analysis, and only the GO categories showing p-values <0.05 in over-representation enrichment analysis were considered further [67]. The assignment of genes into functional categories was manually curated.

### Curcuminoid biosynthesis pathway

Coding sequences of four key genes involved in curcuminoid biosynthesis pathway, namely curcumin synthase 1 (*CURS1*, NCBI accession number BAH56226), curcumin synthase 2 (*CURS2*, NCBI accession number AB506762), curcumin synthase 3 (*CURS3*, NCBI accession number AB506763), and diketide-CoA synthase (*DCS*, NCBI accession number BAH56225) were retrieved [68]. The sequences of these four genes were mapped to the gene set derived from *de novo* transcriptome assembly generated in this study using BLASTN with query coverage ≥50% and e-value 10^-9^ [42]. These sequences were also aligned to *de novo* genome assembly of *Curcuma longa* constructed in this study, using Exonerate v2.2.0 (https://github.com/nathanweeks/exonerate) with 95% of maximal alignment score and 95% quality threshold, and the best hits were selected to construct the gene structures.

## RESULTS

### Sequencing of genome and transcriptome

A total of 94.8 Gbp of 10x Genomics linked read data and 7.7 Gbp of RNA-Seq data was generated from leaf tissue **(Supplementary Tables S1-S2)**. The total genomic data corresponded to ~71X coverage based on the estimated genome size of 1.33 Gbp [26]. To carry out the transcriptome analysis, RNA-Seq data from this study and from the previous studies was used, which together constituted a total of 43.8 Gbp of RNA-Seq data [11], [17], [24], [25], [35]. All the paired-end RNA-Seq reads were trimmed and quality filtered using Trimmomatic v0.38 and used for *de novo* transcriptome assembly. The detailed workflow for genome and transcriptome analysis is shown in **Figure 1**.

**Figure 1.**
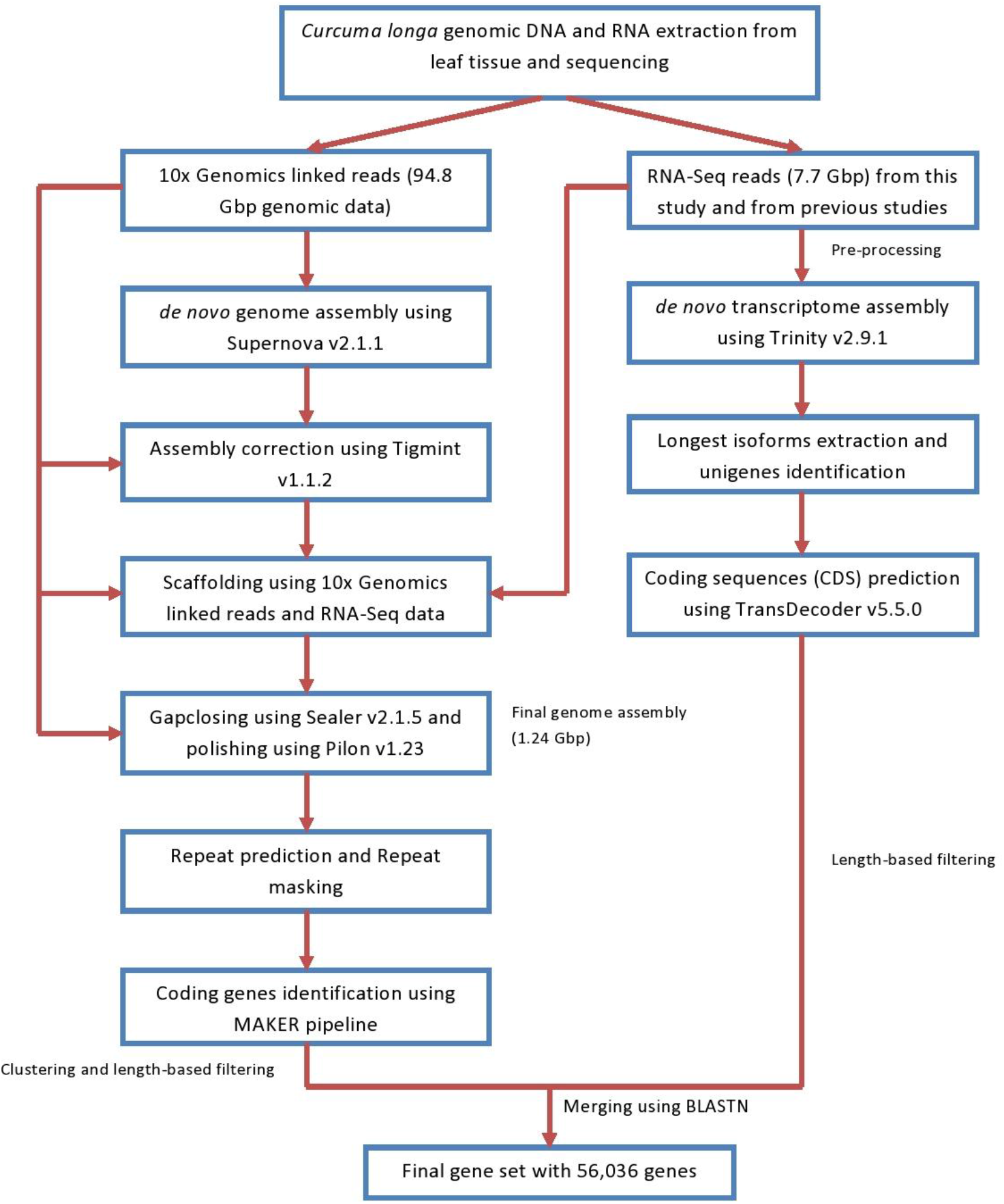
The complete workflow of the genome and transcriptome analysis of *Curcuma longa*

### Assembly of *Curcuma longa* genome and transcriptome

The genome size of *Curcuma longa* was estimated to be 1.15 Gbp using SGA-preqc with barcode-filtered 10x Genomics linked reads, which is close to the previously estimated genome size of 1.33 Gbp [26]. *Curcuma longa* genome assembled using Supernova v2.1.1 had a total size of 1.3 Gbp. After correction of mis-assemblies, scaffolding, gap-closing and polishing, the final draft genome assembly (≥500 bp) of *Curcuma longa* had the total size of 1.24 Gbp that comprised of 1,84,614 scaffolds with N50 value of 18.8 Kbp and GC-content of 38.85% **(Supplementary Table S3)**. BUSCO completeness for Supernova assembled genome was 74% (Complete BUSCOs), which was improved to 81.8% (Complete BUSCOs) in the final polished draft *Curcuma longa* genome assembly **(Supplementary Table S5)**.

The *de novo* transcriptome assembly of *Curcuma longa* using Trinity v2.9.1 had a total size of 3,45,957,265 bp, with a total of 3,63,662 predicted transcripts corresponding to 1,67,582 genes. The complete assembly had an N50 value of 1,605 bp, an average transcript length of 951 bp and GC-content of 42.88% **(Supplementary Table S4).** A total of 1,58,054 unigenes were identified after clustering using CD-HIT-EST v4.8.1 to remove the redundant gene sequences. The coding sequence (CDS) prediction from these unigenes using TransDecoder v5.5.0 resulted in 42,233 coding genes.

### Genome annotation and gene set construction

For repeat identification, a *de novo* custom repeat library was constructed using the final polished *Curcuma longa* genome by RepeatModeler v2.0.1, which resulted in a total of 2,284 repeat families. The repeat families were clustered into 1,821 representative sequences. These were used to soft-mask the genome assembly using RepeatMasker v4.1.0, which predicted 69.92% of *Curcuma longa* genome as repetitive sequences, of which 67.58% was identified as interspersed repeats (30.03% unclassified, 35.41% retroelements and 2.15% DNA transposons). Retroelements consisted of 34.7% LTR repeats (21.04% Ty1/Copia and 13.30% Gypsy/DIRS1 elements) **(Supplementary Table S6)**. Additionally, 6.33% of *Curcuma longa* genome was identified as simple repeats using TRF v4.09. Thus, ~74% of the genome was predicted to be constituted of simple and interspersed repetitive sequences. Among the non-coding RNAs, 3,225 standard amino acid specific tRNAs and 357 hairpin miRNAs were predicted in *Curcuma longa* genome.

MAKER genome annotation pipeline and TransDecoder v5.5.0 predicted a total of 66,396 and 42,233 coding sequences from genome and transcriptome assemblies, respectively. Length-based filtering criteria (≥300 bp) of the above two coding gene sets resulted in 50,744 and 41,785 coding sequences from genome and transcriptome assemblies, respectively. MAKER derived filtered coding sequences were further clustered at 95% sequence identity, which resulted in 40,401 non-redundant coding sequences. These two coding gene sets were merged using BLASTN, resulting in the final *Curcuma longa* gene set comprising of 56,036 coding gene sequences.

### Resolving the phylogenetic position of *Curcuma longa*

From the selected 17 plant species, a total of 1,07,510 orthogroups were identified using protein sequences by OrthoFinder v2.3.9. Among these 1,07,510 orthogroups, 5,294 contained protein sequences from all 17 species, and were used for evolutionary analysis. Further, KinFin v1.0 predicted a total of 1,151 fuzzy one-to-one orthogroups containing protein sequences from all 17 selected species, which were used to construct the maximum likelihood-based phylogenetic tree of *Curcuma longa* with 15 other monocot species and *Arabidopsis thaliana* as an outgroup.

All 1,151 fuzzy one-to-one orthogroups were aligned, concatenated and filtered for undetermined or missing values. This filtered alignment data consisting of 947,855 alignment positions was used to construct the maximum likelihood-based species phylogenetic tree of *Curcuma longa* with 15 other monocot species available on Ensembl plants release 47 and *Arabidopsis thaliana* as the outgroup **(Figure 2).** Position of *Curcuma longa* and the other selected monocots in this genome-wide phylogeny were supported by previously reported phylogenies [69]–[73]. In our phylogeny, *Ananas comosus* showed an earlier divergence among the monocots from Poales order, which was also supported by previously reported studies [69], [72]. Among all selected monocot species in our study, species from Dioscoreales order showed the earliest divergence, supported by the previously reported studies [71], [72]. From our genome-wide phylogeny constructed using the selected monocots, it is apparent that *Musa acuminata* was comparatively closer to *Curcuma longa*. Belonging to the same phylogenetic order Zingiberales, *Curcuma longa* and *Musa acuminata* shared the same clade in the species phylogenetic tree. Species from Zingiberales order are closer to species from Poales order with respect to the species from Dioscoreales order.

**Figure 2.**
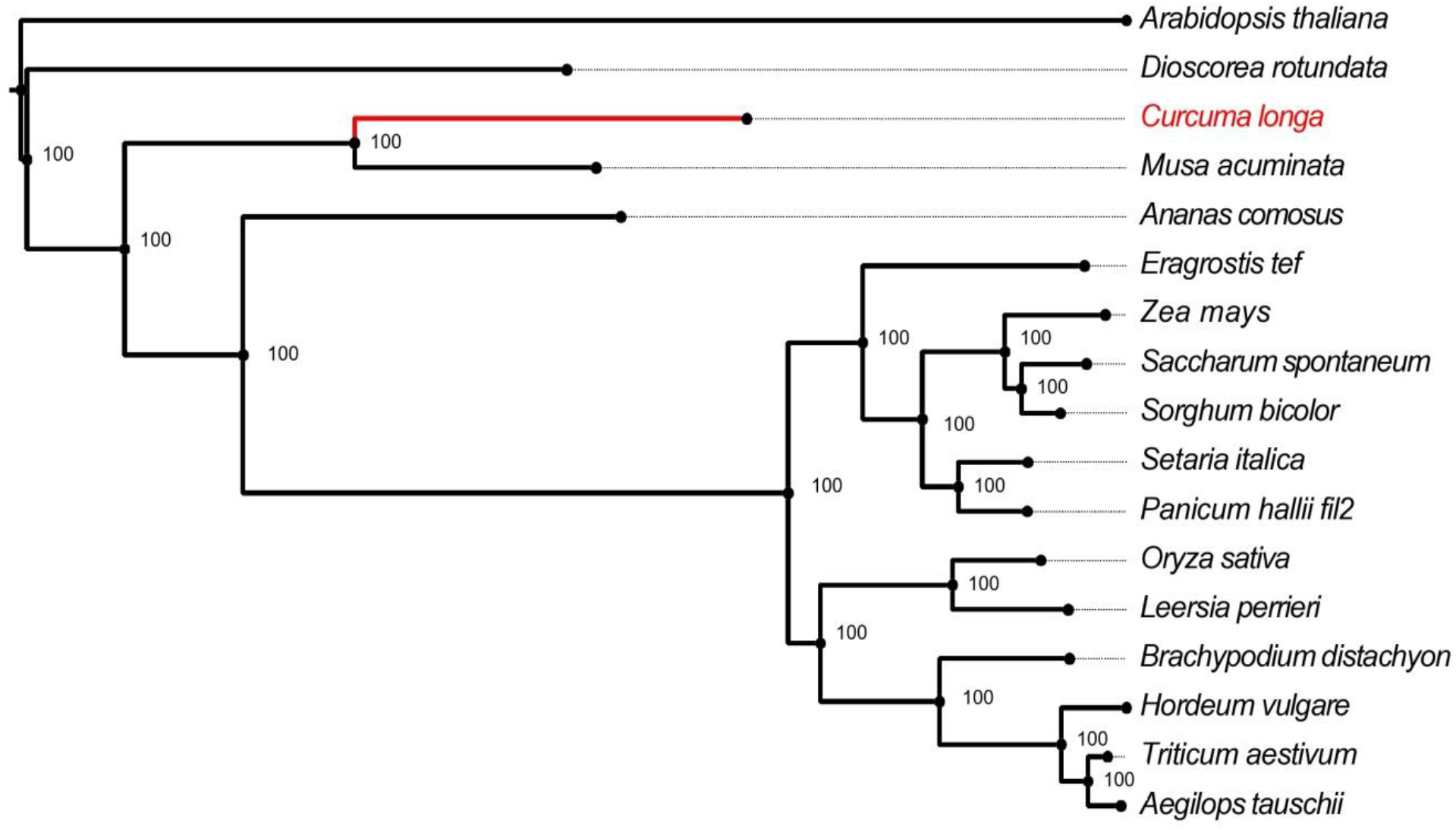
Phylogenetic tree of *Curcuma longa* with 15 other selected species and *Arabidopsis thaliana* as an outgroup species. The values mentioned at the nodes correspond to the bootstrap values.

### Genes with signatures of adaptive evolution

Genes with site-specific signatures of adaptive evolution were identified in *Curcuma longa*. 3,242 genes showed unique amino acid substitution with respect to the other selected species. Among these 3,242 genes, 2,475 genes were identified to have functional impacts using Sorting Intolerant From Tolerant (SIFT), and were considered further. Further, 1,756 genes were found to contain positively selected codon sites with greater than 95% probability. In addition to these site-specific signatures of evolution, 47 genes showed higher rate of nucleotide divergence, and 175 genes showed positive selection in *Curcuma longa* with FDR-corrected p-values <0.05. These positively selected genes had positively selected codon sites with greater than 95% probability.

A total of 109 genes were identified containing more than one of the signatures of adaptive evolution namely positive selection, unique amino acid substitution with functional impact and higher rate of nucleotide divergence. A total of 102 out of 109 MSA genes were associated with categories essential for plant secondary metabolism and defense responses against environmental stress conditions i.e., biotic stress and abiotic stress **(Figure 3)**.

**Figure 3.**
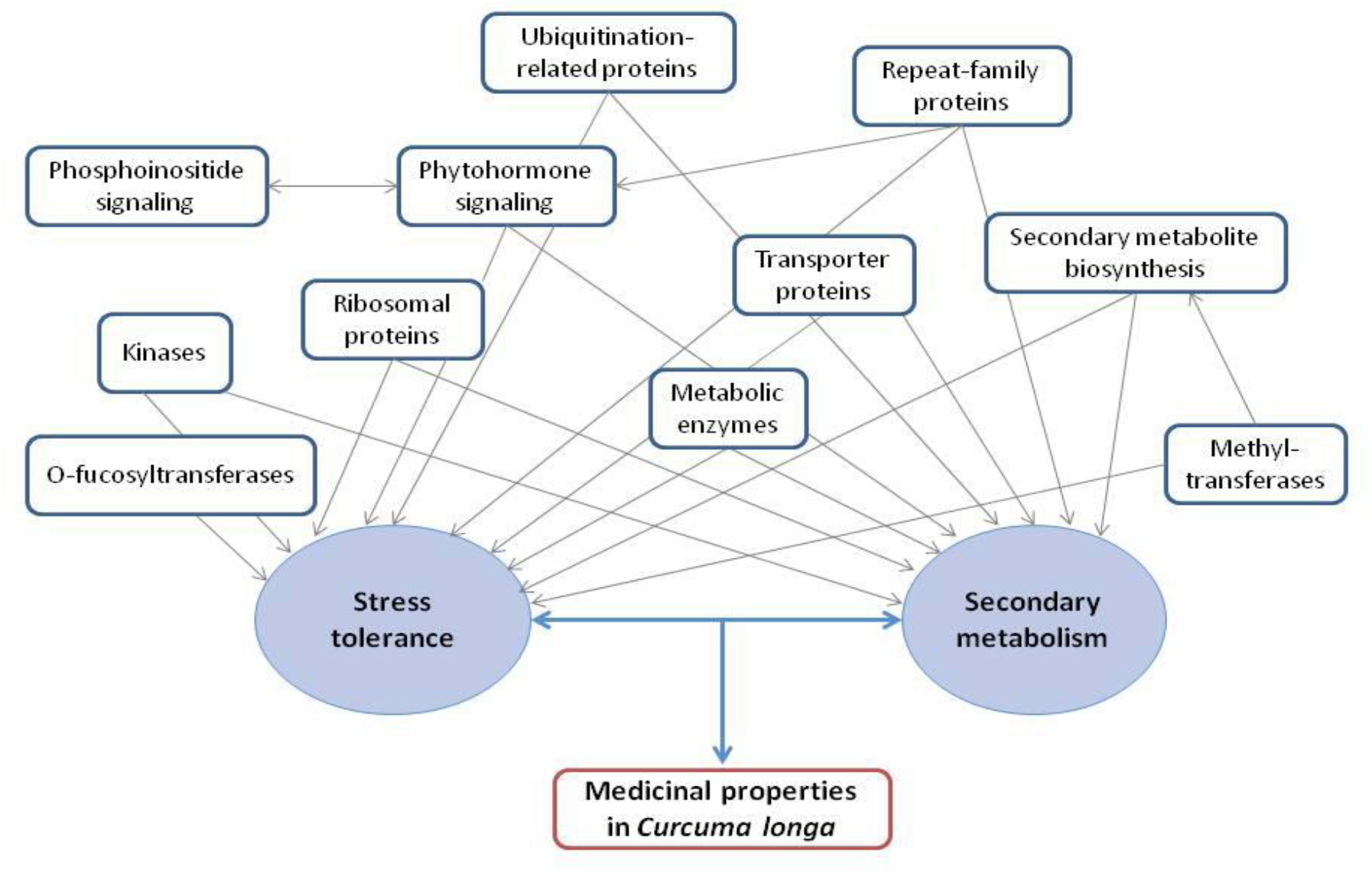
Functional association of MSA genes with medicinal properties of *Curcuma longa*

### Adaptive evolution of plant defense associated genes

Several genes known to provide plant immunity against pathogen infection or disease were found in MSA genes. Since plants lack adaptive immune response, the innate immune response in plants is provided by PAMP-triggered (PTI) and effector-triggered (ETI) immunity with the help of two plant stress hormones, salicylic acid (SA) and jasmonic acid (JA) signaling pathway [74]. Among these MSA genes, *‘IAR3’* is important for production of 12-Hydroxyjasmonic acid, *‘LOX’* gene helps in oxylipins such as JA production by oxidation of fatty acids, *‘LOB’* gene regulates anthocyanin metabolism and components of JA signaling pathways and is modulated by methyl jasmonate treatment, *‘AT4G32690’* regulates nitric oxide mediated JA signaling and JA accumulation in plants, and *AOC3’* gene gets upregulated by methyl jasmonate treatment [75]–[79]. Also, *‘WRKY22’* transcription factor is induced by pathogen attack and is involved in SA-signaling pathway mediated plant immunity, *‘PAT14’* gene negatively regulates SA-mediated defense response, and L,L-diaminopimelate aminotransferase negatively regulates SA levels and PR gene expression [74], [80], [81]. Jasmonic acid and salicylic acid elicit the production and accumulation of secondary metabolites such as phenolics, terpenoids, alkaloids, and glycosides, in medicinal plants [82].

Three O-fucosyltransferase family proteins – *‘AT1G76270’, ‘AT1G62330’*, and *‘AT1G04910’* were found in MSA genes category, and are involved in plant immunity. Previously it has been shown that the lack of fucosylation of genes led to increased disease susceptibility in *Arabidopsis sp*. by affecting PTI, ETI as well as stomatal and apoplastic defense [83]. Ubiquitin-conjugating enzyme *‘AT2G18600’* and three E3 ubiquitin-protein ligase genes – *‘AT2G17020’, ‘AT5G04460’, ‘SPL2’* also showed multiple signs of adaptive evolution. Ubiquitination-related proteins affect hypersensitive-response (HR) and phytohormone signaling mediated pathogen defense by targeting proteins for proteasomal degradation [84]–[87]. These ubiquitin ligase proteins also regulate various abiotic stress responses e.g., drought, salinity, temperature [87]. Among the MSA genes, six ribosomal subunit proteins are regulated by signaling molecules such as methyl jasmonate, salicylic acid and environmental stress e.g., cold, heat, UV, drought, salinity [88], [89]. Two cell cycle related MSA genes – cyclin-A and *‘APC10’* are involved in plant immunity, disease resistance as well as abiotic stress responses [90], [91].

Three genes related to phosphoinositide signaling were found to be among the MSA genes. Phosphoinositide signaling, which is modulated in response to mechanical wounding in *Arabidopsis thaliana*, contribute to phytohormones mediated defense responses [92]. This signaling cascade also plays a role in abiotic stress response such as water stress [93]. Two *ERD* family proteins – *‘AT4G02900’* and *‘AT1G30360’* were identified as MSA genes. *ERD* genes are known to provide abiotic stress tolerance via drought tolerance and salicylic acid-dependent plant defense [94]. Six different repeat family genes belonging to WD-family repeats, TPR-like superfamily repeats, ARM superfamily repeats and Galactose oxidase/kelch superfamily repeats were identified as MSA genes. These repeat family proteins are involved in plant innate immunity as well as abiotic stress tolerance [95]–[98].

### Abundance of secondary metabolism pathways in *Curcuma longa* genome

Genes associated with secondary metabolite biosynthesis and secondary metabolism were found to be abundant (64 out of 109 genes) among the MSA genes in *Curcuma longa*. Of these, *‘DWF4’* is involved in brassinosteroid biosynthesis, *‘PGT1’* has a role in phenylpropanoid biosynthesis, *‘AT4G14420’* aids in glucosinolate biosynthesis [99]–[101]. Methyltransferases e.g., *‘AT1G3317ff*, *‘OMT’, ‘AT1G50000’* are also involved in secondary metabolites biosynthesis and responsible for methylation of secondary metabolites, which in turn assist in disease resistance and stress tolerance in plants [102], [103]. These secondary metabolites possess various medicinal applications such as anti-inflammatory, antimicrobial, antioxidant, and anti-carcinogenic properties [7], [16], [104], [105]. Transporter genes e.g., *‘MRS2’* and *‘AT1G29830* magnesium transporters, *NRT* a nitrate or nitrite transporter, *‘IREG3’* an iron-regulated transporter, *TC.APA’* a polyamine transporter, and *‘AT1G51610’* a cation efflux family protein also showed multiple signs of evolution. The evolution of these transporter genes appears significant since the alteration in concentration of magnesium and iron metal ions regulate accumulation and translocation of secondary metabolites [106]–[110].

Polyketides, such as curcuminoids, represent a diverse class of secondary metabolites, which are crucial for plants survival under environmental challenges [111]. Curcuminoid biosynthesis pathway is the most important secondary metabolism pathway in *C. longa* where phenylalanine is converted to coumaroyl-CoA via cinnamic acid and coumaric acid [112]. Notably, the *‘OMT* gene, which is involved in conversion of coumaroyl-CoA to feruloyl-CoA in the curcuminoid biosynthesis pathway [113], showed positive selection and unique amino acid substitution in this study.

Further, coumaroyl-CoA and feruloyl-CoA are used for the production of curcumin, demethoxycurcumin, and bisdemethoxycurcumin via coumaroyl-diketide-CoA and feruloyl-diketide-CoA, catalyzed by four enzymes - *‘CURS1’, ‘CURS2’, ‘CURS3’*, and *‘DCS’* [112]. To identify these enzymes in the genome and transcriptome assemblies constructed in this study, we mapped the coding gene sequences of *‘CURS1’, ‘CURS2’, ‘CURS3’*, and *‘DCS’* genes on these assemblies. All four enzymes were found to be present in the *de novo* genome assembly and in the gene set derived from *de novo* transcriptome assembly of *Curcuma longa* **(Supplementary Table S7).** Using Exonerate tool, we further constructed the gene structures for these four major curcuminoid biosynthesis genes. Each of the *‘CURS1’, ‘CURS2’* and *‘CURS3’* genes consisted of two exons and one intron. *‘DCS’* gene consisted of three exons and two introns. *‘DCS’* and *‘CURS’* genes are members of chalcone synthase (*CHS*) family [114], and the genes from *CHS* family generally consist of two exons and one intron [115], which is consistent with and also further supported by our findings.

## DISCUSSION

*Curcuma longa* is a monocot species from Zingiberaceae plant family and is widely known for its medicinal properties and therapeutic applications [19], [116]. In this study, we carried out the whole genome sequencing and reported the draft genome sequence of *Curcuma longa*. This is the first genome sequenced from Zingiberaceae plant family, which comprises more than 1300 species, and thus will act as a valuable reference for studying the members of this family including those of *Curcuma* genus. The application of 10x Genomics linked read sequencing that has the potential to resolve complex polyploid genomes [117], helped in successfully constructing the *Curcuma longa* draft genome of 1.24 Gbp with a decent N50 of 18.8 Kbp. After assembly correction, scaffolding, gap-closing and polishing, the BUSCO completeness of Supernova v2.1.1 derived *Curcuma longa* genome improved to 81.8%, which is similar to other plant genomes, thus indicating the usefulness of post-assembly processing [118].

Since the construction of a comprehensive gene set was essential to explore the genetic basis of its medicinal properties, both genome and transcriptome assemblies, and an integrated approach using *de novo* and homology-based methods were used resulting in the final set of 56,036 genes. The identification of all Type III polyketide synthase genes *‘CURS1’, ‘CURS2’, ‘CURS3’*, and *‘DCS’*, involved in the biosynthesis of the three most important secondary metabolites (curcuminoids) – curcumin, demethoxycurcumin and bisdemethoxycurcumin, in both genome and transcriptome assemblies attests to the quality and comprehensiveness of our genome and transcriptome assembly. Further, the revelation of complete gene structures of the above four biosynthesis genes of curcuminoid pathway from the draft genome of this plant is likely to help further studies and for better commercial exploitation of these curcuminoids that find wide applications as coloring agents, food additives and possess antioxidant, anti-inflammatory, anti-microbial, neuroprotective, anti-cancer and many other medicinal properties [119].

Repetitive sequence prediction revealed that ~74% of the genome consisted of repeat elements, which is similar to other plant genomes [120]. Notably among the LTR repeat elements, Ty1/Copia elements (21.04%) were more abundant than Gypsy/DIRS1 elements (13.30%), which corroborates with the observations made in the case of *Musa acuminata* species from the same Zingiberales plant order, and thus appears to be a specific signature of repeat elements in Zingiberales order [121].

The genome-wide phylogenetic analysis of *Curcuma longa* with 15 other representative monocot species available on Ensembl plants revealed the relative position of *Curcuma longa*, which was supported by previously reported phylogenies using 1,685 gene partitions, and using phytocystatin gene *‘CypCl’* [69], [70]. Ren et al. (2018) also showed similar phylogenetic position of *Curcuma longa* with other selected monocots – *Dioscorea sp., Musa acuminata, Brachypodium distachyon, Oryza sativa, Panicum sp., Setaria italica, Zea mays, Sorghum bicolor* using genome and transcriptome data of 105 angiosperms [71]. Also, an updated megaphylogeny for vascular plants showed similar relative phylogenetic position of *Curcuma longa* with respect to the selected monocots [72]. Further, selected species from Poales order also showed similar relative positions with respect to each other [72], [73]. Absence of any polytomy in the phylogenetic tree is because of large number of genomic loci and a high bootstrap value, or no multiple speciation events took place at the same time. Taken together, the genome-wide phylogenetic analysis of *Curcuma longa* confirmed its phylogenetic position and will be a useful reference for further studies.

Analysis of genes with signatures of adaptive evolution using 5,294 orthologous gene sets revealed that a large proportion (~94%) of the genes with multiple signs of adaptive evolution (MSA) were associated with plant defense mechanisms against biotic and abiotic stress responses, and secondary metabolism. Notable ones among these are the genes associated with Jasmonic acid and salicylic acid signaling pathways. These two pathways are an important components of plant innate immune response [74], and also affect plant secondary metabolism by regulating the production of secondary metabolites [82], thus play a crucial role in plant-pathogen interaction. Jasmonic acid is also reported to have a role in induction and growth of rhizome in vitro through its interaction with ethylene, which is important for a rhizomatous plant like *C. longa* [122]. Further, one of the genes (‘*OMT’* gene) for the enzymes involved in curcuminoid biosynthesis pathway was also found to have signatures of adaptive evolution, which is an important observation because curcuminoid is the most important secondary metabolite of *C. longa*. The genes for the other four key enzymes *(CURS1, CURS2, CURS3, DCS)* of this pathway could not be found in the list of MSA genes since these genes are unique to *Curcuma* genus and were absent in the other species considered for the evolutionary analysis. It is important to mention here that the biosynthesis of secondary metabolites such as polyketides (curcuminoids), which are crucial for plants survival under environmental challenges, are regulated by biotic and abiotic stress responses [123]–[125]. Also, it is known that secondary metabolites in plants are primarily produced in response to environmental stress and for plant defense for better survival under various environmental conditions [126], and several of these secondary metabolites also possess medicinal value. This also seems to be the case with *C. longa* where the abundance of adaptively evolved genes associated with plant defense mechanisms and secondary metabolism makes it tempting to speculate that these genes gradually evolved to confer resistance and environmental adaptation for a perennial rhizomatous plant like *C. longa*. Further, several of the metabolites produced in the above processes for conferring resistance and environmental adaptation to *C. longa* possess diverse medicinal properties, and provides turmeric with its medicinal characteristics and traditional significance.

## CONCLUSION

Due to the enormous medicinal and traditional value of *Curcuma longa*, the revelation of its genome sequence was much needed and was achieved in this study. Further, this genome is the first genome sequenced so far from the Zingiberaceae plant family. The genomic analysis revealed multiple signs of adaptive evolution in genes associated with plant defense responses against biotic and abiotic challenges and secondary metabolism, which perhaps are evolved for its perennial persistence and also forms the basis of its medicinal properties. Thus, the availability of draft genome and transcriptome, gene set and associated genomic information, and the comprehensive evolutionary analysis with other monocot species will be a valuable reference to gain further insights into the evolution of diverse medicinal properties of this plant and other members of Zingiberaceae plant family.

## LIST OF ABBREVIATIONS

PCR: Polymerase Chain Reaction
BLAST: Basic Local Alignment Search Tool
miRNA: micro RNA
tRNA: transfer RNA
ORF: Open Reading Frame
LTR: Long Terminal Repeat
FDR: False discovery rate
MSA: Multiple signs of adaptive evolution
*DWF4*: Dwarf 4
*PGT1*: Phloretin 2’-O-glucosyltransferase
*OMT*: O-methyltransferase
*MRS2*: Mitochondrial RNA Splicing 2
*NRT*: Nitrate Transporter
*IREG3*: Iron Regulated 3
*TC.APA*: basic amino acid/polyamine antiporter
*IAR3*: IAA-Alanine Resistant 3
*LOX*: Lipoxygenase
*LOB*: LATERAL ORGAN BOUNDARIES
*AOC3*: Amine Oxidase Copper Containing 3
*PAT14*: Protein Acyltransferase 14
PR: Pathogenesis-related
PTI: PAMP-triggered immunity
ETI: Effector-triggered immunity
*SPL2*: SP1-Like 2
UV: Ultraviolet
*APC10*: Anaphase Promoting Complex subunit 10
*ERD*: Early-Responsive to Dehydration stress
TPR: Tetratrico Peptide Repeat
ARM: Armadillo
BUSCO: Benchmarking Universal Single-Copy Orthologs
KEGG: Kyoto Encyclopedia of Genes and Genomes
GO: Gene ontology
COG: Clusters of Orthologous Groups
*DCS*: diketide-CoA synthase
*CURS1*: Curcumin synthase 1
*CURS2*: Curcumin synthase 2
*CURS3*: Curcumin synthase 3

## COMPETING INTERESTS

The authors declare no competing financial and non-financial interest.

## AUTHORS’ CONTRIBUTIONS

VKS conceived and coordinated the project. SM prepared the DNA and RNA samples, prepared the samples for sequencing, and performed the species identification assay. AC and VKS designed the computational framework of the study. AC performed all the computational analysis presented in the study. AC, SM, and SKJ performed the functional annotation of gene sets. AC, SM, SKJ and VKS analysed the data and interpreted the results. AC constructed the figures. AC, SM, SKJ and VKS wrote the manuscript. All the authors have read and approved the final version of the manuscript.

## ACKNOWLEDGEMENTS

AC and SM thank Council of Scientific and Industrial Research (CSIR) for fellowship. SKJ thanks Department of Science and Technology for the DST-INSPIRE fellowship. The authors thank the intramural research funds provided by IISER Bhopal.

## DATA AVAILABILITY

The raw genome and transcriptome reads of *Curcuma longa* have been deposited in National Center for Biotechnology Information (NCBI).

## Notes

### Competing Interest Statement

The authors have declared no competing interest.

