## Supplementary Material for "Genome sequencing of turmeric provides evolutionary insights into its medicinal properties"

**SUPPLEMENTARY TABLES**

**Supplementary Table S1. Summary of the 10x Genomics linked read sequence data for *Curcuma longa* genome**

| Average Read Length | Number of Reads | Total Data | Sequencing data coverage |
|---|---|---|---|
| 150 bp | 631.11 million | 94.8 Gb | ~71x |

The sequencing coverage was calculated with respect to the estimated genome size of 1.33 Gb [1].

**Supplementary Table S2. Summary of RNA-Seq data for *Curcuma longa* transcriptome**

| Tissue | Average Read Length R1 (bp) | Average Read Length R2 (bp) | Total Number of Read pairs | Total Number of Bases in R1 (bp) | Total Number of Bases in R2 (bp) | Total Number of Bases (bp) |
|---|---|---|---|---|---|---|
| Leaf | 133.8 | 133.8 | 28,960,676 | 3,874,053,529 | 3,874,120,943 | 7,748,174,472 |

In addition to our data, RNA-seq data from previously reported studies were used, which resulted in the total transcriptome data of 43.8 Gb [2]–[6].

**Supplementary Table S3. Summary statistics of final *de novo* draft genome assembly of *Curcuma longa* (≥500 bp) using 10x Genomics linked reads**

| Parameter | Value |
|---|---|
| Number of contigs (>= 500 bp) | 1,84,614 |
| Number of contigs (>= 1000 bp) | 1,34,864 |
| Number of contigs (>= 5000 bp) | 56,983 |
| Number of contigs (>= 10000 bp) | 28,243 |
| Number of contigs (>= 25000 bp) | 9,325 |
| Number of contigs (>= 50000 bp) | 3,285 |
| Total length (>= 500 bp) | 1,242,536,276 |
| Total length (>= 1000 bp) | 1,206,928,265 |
| Total length (>= 5000 bp) | 1,019,273,423 |
| Total length (>= 10000 bp) | 818,136,386 |
| Total length (>= 25000 bp) | 525,670,954 |

| | |
|---|---|
| Total length (>= 50000 bp) | 319,823,365 |
| Largest contig (bp) | 706,167 |
| GC (%) | 38.85 |
| N50 | 18,798 |
| L50 | 13,753 |
| Number of N's per 100 kbp | 2111.22 |

**Supplementary Table S4. Summary statistics of *de novo* transcriptome assembly of *Curcuma longa***

| **Statistics based on all transcript contigs** | |
|---|---|
| Contig N50 | 1,605 |
| Median contig length | 549 |
| Average contig | 951.32 |
| Total assembled bases | 345,957,265 |
| **Statistics based on only longest isoform per gene** | |
| Contig N50 | 1,060 |
| Median contig length | 356 |
| Average contig | 663.57 |
| Total assembled bases | 111,201,747 |
| **Counts of genes and transcripts** | |
| Total trinity 'genes' | 1,67,582 |
| Total trinity transcripts | 3,63,662 |
| GC (%) | 42.88 |

**Supplementary Table S5. BUSCO statistics of *Curcuma longa* genome**

| **Parameters** | **Supernova v2.1.1 assembled raw genome** | **Final *Curcuma longa* draft genome (≥500 bp)** |
|---|---|---|
| Complete BUSCOs (C) | 1,195 (74.0%) | 1,319 (81.8%) |
| Fragmented BUSCOs (F) | 203 (12.6%) | 130 (8.1%) |
| Missing BUSCOs (M) | 216 (13.4%) | 165 (10.1%) |

| | | |
|---|---|---|
| Total BUSCO groups searched | 1,614 | 1,614 |

BUSCO analysis was performed using the reference BUSCO database embryophyta_odb10.

**Supplementary Table S6. Summary statistics of repetitive sequences in *Curcuma longa* genome from RepeatMasker**

| | | | | | |
|---|---|---|---|---|---|
| Number of sequences: | 1,84,614 | | | | |
| Total length (bp): | 1,242,536,276 bp | | | | |
| GC (%) | 38.85 % | | | | |
| Bases masked: | 868,821,943 bp ( 69.92 %) | | | | |
| | | | Number of Elements | Length occupied (bp) | Percentage of Sequence |
| Retroelements | | | 3,45,843 | 440,028,208 bp | 35.41 % |
| | SINEs | | 2,132 | 25,36,945 bp | 0.20 % |
| | Penelope | | 0 | 0 bp | 0.00% |
| | LINEs | | 13,021 | 63,52,436 bp | 0.51 % |
| | | CRE/SLACS | 0 | 0 bp | 0.00% |
| | | L2/CR1/Rex | 0 | 0 bp | 0.00% |
| | | R1/LOA/Jockey | 2,050 | 26,65,712 bp | 0.21 % |
| | | R2/R4/NeSL | 0 | 0 bp | 0.00% |
| | | RTE/Bov-B | 7,326 | 15,29,090 bp | 0.12 % |
| | | L1/CIN4 | 3,645 | 21,57,634 bp | 0.17 % |
| | LTR elements: | | 3,30,690 | 431,138,827 bp | 34.70 % |
| | | BEL/Pao | 0 | 0 bp | 0.00% |
| | | Ty1/Copia | 1,98,955 | 261,440,475 bp | 21.04 % |
| | | Gypsy/DIRS1 | 1,22,625 | 165,316,610 bp | 13.30 % |
| | | Retroviral | 4,000 | 10,17,184 bp | 0.08 % |

| | | | | | |
|---|---|---|---|---|---|
| DNA transposons | | | 34,307 | 2,66,64,558 bp | 2.15 % |
| | hobo-Activator | | 11,999 | 58,59,232 bp | 0.47 % |
| | Tc1-IS630-Pogo | | 0 | 0 bp | 0.00% |
| | En-Spm | | 0 | 0 bp | 0.00% |
| | MuDR-IS905 | | 0 | 0 bp | 0.00% |
| | PiggyBac | | 0 | 0 bp | 0.00% |
| | Tourist/Harbinger | | 2,624 | 13,74,155 bp | 0.11 % |
| | Other (Mirage, P-element, Transib) | | 0 | 0 bp | 0.00% |
| Rolling-circles | | | 27,802 | 1,61,85,798 bp | 1.30 % |
| Unclassified: | | | 10,34,622 | 373,075,264 bp | 30.03 % |
| Total interspersed repeats: | | | | 839,768,030 bp | 67.58 % |
| Small RNA: | | | 5,788 | 52,96,422 bp | 0.43 % |
| Satellites: | | | 0 | 0 bp | 0.00% |
| Simple repeats: | | | 1,39,683 | 62,11,866 bp | 0.50 % |
| Low complexity: | | | 26,274 | 13,59,827 bp | 0.11 % |

Note: The information provided in the above table is as per the default output of RepeatMasker program.

**Supplementary Table S7. Summary of curcuminoid biosynthesis genes mapping on the gene set derived from *de novo* transcriptome assembly**

| Query gene | Query length | Percentage of identical matches | Alignment length (bp) | e-value |
|---|---|---|---|---|
| curcumin synthase 1 | 1170 bp | 83.333 | 876 | 0.0 |
| curcumin synthase 2 | 1176 bp | 99.220 | 897 | 0.0 |
| curcumin synthase 3 | 1173 bp | 99.420 | 690 | 0.0 |
| diketide-CoA synthase | 1170 bp | 88.761 | 1130 | 0.0 |

Mapping statistics was calculated based on the best hits obtained from BLASTN search with query coverage ≥50% and e-value $10^{-9}$.

### SUPPLEMENTARY TEXTS

#### Supplementary Text S1.

**Sample collection and DNA Extraction**

The plant sample was collected from an agricultural farm located in Bhopal, India (23.2280252°N 77.2088987°E). The plant was brought to lab and processed immediately. The leaves were used for DNA extraction using Carlson buffer [100mM Tris HCl, 2% CTAB (Cetyl Trimethyl Ammonium Bromide), 1.4M NaCl, 1% PEG 8000, 20 mM EDTA pH 9.5] supplemented with β-mercaptoethanol (2.5 µl for each ml of buffer) for lysis [7]. Before adding the sample to buffer, it was allowed to pre-heat at 65˚C for 30 mins. The homogenized powder of leaves was added to the pre-heated buffer (1 ml) which was supplemented with 2 µl of RNase A (20 mg/ml) and 25 µl of Proteinase K (20 µL/mL). The tubes were given 1 hour incubation at 65˚C with in-between mixing. In order to obtain high molecular DNA the tubes were mixed by inverting the tubes in all the steps. All the centrifugation steps were performed at 5,000xg at 4˚C. After cooling the tubes at room temperature, 1 ml of chloroform was added and centrifuged for 15 mins. The aqueous layer formed was transferred to new centrifuge tube, ice-cold Isopropanol (0.7x volume) was added and followed by an overnight incubation at -20˚C in order to facilitate DNA precipitation. The tubes were centrifuged for 45 mins to pellet down the precipitated DNA. The DNA pellet was dissolved in

500 µl of G2 buffer (QiaAmp Blood and Cell culture Kit) by incubating at 50˚C for 15 mins. The Genomic tip 20 was equilibrated and the dissolved DNA was allowed to pass through it. Here, multiple tubes (up to 2 ml of G2 buffer) were loaded to single genomic tip 20 in order to increase the yield. Buffers were allowed to pass through the Genomic tip 20 via gravity flow. 1 ml of QC buffer was used thrice for washing the column and then DNA elution was done in 1 ml of QF buffer (pre-heated at 55˚C). The DNA was precipitated with 0.7X volume of ice-cold isopropanol and facilitated by an overnight incubation at -20˚C. The DNA was centrifuged for 30 mins at 4˚C to pellet it down. The DNA pellet was washed with 1 ml of ice-cold 70 % ethanol, air dried to remove the residual ethanol and finally eluted in 50 µl of nuclease free water. The NanoDrop$^{TM}$8000 Spectrophotometer (ThermoFisherScientific, USA) and 0.8-1% agarose gel electrophoresis, and Qubit 2.0 Fluorometer using Qubit dsDNA BR assay kit (Invitrogen, USA) were used to assess the quality and quantity of extracted DNA, respectively.

**Species Identification**

Species identification was done by using primers for a nuclear gene (Internal Transcribed Spacer ITS) and a chloroplast gene (Maturase K). The forward and reverse primers for complete ITS amplification were 5′-TCCGTAGGTGAACCTGCGG-3′ and 5′-TCCTCCGCTTATTGATATGC-3′, respectively. For ITS 2 gene amplification the primer set used was 5′-GCATCGATGAAGAACGCAGC-3′ and 5′-TCCTCCGCTTATTGATATGC-3′ as forward and reverse primers, respectively. The PCR programme ran on Veriti 96 well thermal cycler (Applied Biosystems) for ITS gene amplification was 94 ˚C for 3 mins, 35 cycles of 94 ˚C for 1 min, 55 ˚C for 1 min and 72 ˚C for 2.5 mins and 72 ˚C for 10 mins. Similarly, the primer set for Maturase K (MatK) included 5′-CGATCTATTCATTCAATATTTC-3′ as forward primer and 5′-TCTAGCACACGAAAGTCGAAGT-3′ as reverse primer. The PCR programme ran for MatK was 95 ˚C for 3 mins, 35 cycles of 95 ˚C for 30 sec, 50 ˚C for 3 mins and 72 ˚C for 1:15 min and final extension at 72 ˚C for 7 mins. The amplification was assessed on 2X agarose gel electrophoresis. The PCR product was purified and sequenced at in-house Sanger sequencing facility. The species was confirmed as *Curcuma longa* by checking the sequence identity and alignment with NCBI database using BLASTN.

**Transcriptome Extraction**

For RNA extraction, the powdered leaves (50-100mg) were added to 1 ml of TriZol reagent (Invitrogen, USA) and shaken for 5 mins. For ensuring complete dissociation of nucleoprotein complexes, the tubes were incubated at room temperature for 5 mins. To each tube, 200 ul of chloroform was added and vortexed for 15 seconds followed by incubation of 10 mins at room temperature (RT). In order to separate phases, the tubes were centrifuged at 12,000xg for 15 mins at 4˚C. The upper aqueous layer was transferred to a new tube where the RNA was allowed to precipitate with 500 ul of ice-cold isopropanol by incubating at RT for 5-10 mins. The RNA was pelleted by centrifuging at 12,000xg for 10 mins at 4˚C. Washing of RNA pellet was done with 1 ml of 75% ethanol. The pellet was dried by keeping it at 37˚C for 30 mins. The RNA pellet was resuspended in 30ul of nuclease free water by incubating it at 55-60˚C for 10-15 mins [8]. The quality and quantity of RNA was assessed by NanoDrop™8000 Spectrophotometer (ThermoFisherScientific, USA) and Qubit 2.0 Fluorometer using Qubit ssRNA HS assay kit (Invitrogen, USA), respectively.

**Genomic and Transcriptomic Sequencing**

For genomic sequencing the DNA library was prepared using Chromium Controller instrument, Chromium™ Genome Library & Gel Bead Kit v2 (10x Genomics) by following the manufacturer's instructions. The transcriptomic library was prepared with TruSeq Stranded Total RNA Library Preparation kit (Illumina, Inc., United States) by following the manufacturer's protocol with Ribo-Zero workflow. The quality of libraries was evaluated on Agilent 2200 TapeStation using High Sensitivity D1000 ScreenTape (Agilent, Santa Clara, CA). Both the libraries (Genomic and transcriptomic) were sequenced on Novaseq 6000 (Illumina, Inc., United States) generating 150 base pair paired end reads.

**Supplementary Text S2.**

**Genome size estimation**

Barcode sequences were trimmed from raw 10x Genomics linked reads using a set of python scripts (https://github.com/ucdavis-bioinformatics/proc10xG). Barcodes were extracted from all paired-end linked reads using process_10xReads.py script with default parameters and irrespective of presence of valid gem barcodes. Reads were then filtered based on barcode status, using filter_10xReads.py script.

Genome size was estimated using a k-mer count distribution method implemented in SGA-preqc, that removes error prone k-mers with low occurrence count [9] . First, sga preprocess was used in paired-end mode with filtered linked reads; then the preprocessed reads were indexed with 'ropebwt' indexing algorithm and '--no-reverse' option; finally sga preqc was run with default settings to estimate the genome size.

**Genome assembly and polishing**

A total of 631.11 million 10x Genomics linked reads, corresponding to ~71x sequencing coverage, was used without any pre-processing, for *de novo* assembly of *Curcuma longa* genome using Supernova v2.1.1 with maxreads=all option and Supernova mkoutput in 'pseudohap' style was used to generate the haplotype-phased fasta assembly file [10]. Barcodes from the raw reads were processed using Longranger basic v2.2.2 (https://support.10xgenomics.com/genome-exome/software/pipelines/latest/installation), to use in further assembly post-processing purpose. Tigmint v1.1.2 was used for correcting the mis-assemblies using the long range information resided within the linked-reads [11]. First, the assembled genome was indexed and barcode-processed linked reads were mapped using BWA-MEM and samtools v1.9 was used to generate the ".bam" file [12], [13]. This ".bam" file was used by tigmint-molecule to generate the ".bed" file that is further used by tigmint-cut to cut the mis-assembled regions and generate the corrected assembly.

For first round of scaffolding, linked reads were mapped to the corrected genome assembly using Longranger align v2.2.2 (https://support.10xgenomics.com/genome-exome/software/pipelines/latest/installation) and samtools v1.9 was used to generate the ".bam" file [13]. Using this ".bam" file, combination of ARCS v1.1.1 and LINKS v1.8.6 was used with default parameters, to generate a more contiguous assembly [14], [15]. For further scaffolding, AGOUTI v0.3.3 was used with the filtered paired-end RNA-Seq reads that were required for *de novo* transcriptome assembly [16]. These paired-end RNA-Seq reads were mapped to previously scaffolded genome using BWA-MEM and samtools v1.9 was used to generate the ".bam" file [12], [13]. Using this ".bam" file and AUGUSTUS v3.2.3 derived ".gff3" file [17], AGOUTI was used to generate further scaffolded genome assembly [16].

Sealer v2.1.5 was used to gap-close this scaffolded genome assembly with barcode-processed linked reads and k-mer values from 30 to 120 (with an interval of 10 bp) with a Bloom-filter size of 950 GB [18]. Barcode-processed linked reads were again mapped to this gap-closed genome assembly using BWA-MEM and samtools v1.9 was used to generate the ".bam" file [12], [13]. Finally, Pilon v1.23 was used with this ".bam" file to polish the genome assembly and improve the assembly quality [19].

**Transcriptome data pre-processing**

Trimmomatic v0.38 was used for pre-processing of RNA-Seq data [20]. Adapter trimming was performed with maximum 2 mismatches in a 16-base seed matching; 30 and 10 was used as palindrome clip threshold and simple clip threshold, respectively. Quality threshold was set to 20 for leading and trailing end trimming of a read. Sliding window trimming for the reads was performed if an average quality score for a 4-base window went under 20. All reads containing less than 60 bases were removed.

**Supplementary Text S3.**

**Tandem Repeats identification**

For tandem repeat identification on the final polished *Curcuma longa* draft genome (≥500 bp), Tandem Repeat Finder (TRF) v4.09 was used with the parameters as follows: matching weight = 2, mismatching penalty = 7, indel penalty = 7, match probability = 80%, indel probability = 10%, minimum alignment score = 50, and maximum period size = 2000 [21].

**Identification of transfer RNAs (tRNAs)**

tRNAscan-SE v2.0.5 was used for *de novo* prediction of tRNAs in final polished *Curcuma longa* draft genome assembly (≥500 bp) with default parameters [22]. A total of 3,462 tRNAs were predicted, which were further classified as follows:

tRNAs decoding Standard 20 AA: 3,225

Selenocysteine tRNAs (TCA): 0

Possible suppressor tRNAs (CTA,TTA,TCA): 2

tRNAs with undetermined/unknown isotypes: 24

Predicted pseudogenes: 211

Along with these, 107 tRNAs with introns were identified.

**Identification of microRNAs**

miRBase database was used for homology-based identification of hairpin miRNAs [23]. A total of 38,589 hairpin miRNAs were clustered using CD-HIT-EST v4.8.1 with 90% sequence identity, to generate 22,365 non-redundant sequences [24]. Using these sequences, BLASTN was used with parameters of 80% identity and e-value 1e-03, to identify the hairpin miRNAs in *Curcuma longa* final draft genome assembly [25].

[11] S. D. Jackman *et al.*, “Tigmint: Correcting assembly errors using linked reads from large

molecules," *BMC Bioinformatics*, 2018, doi: 10.1186/s12859-018-2425-6.

[22] P. P. Chan and T. M. Lowe, "tRNAscan-SE: Searching for tRNA genes in genomic sequences," in *Methods in Molecular Biology*, 2019.

[23] S. Griffiths-Jones, H. K. Saini, S. Van Dongen, and A. J. Enright, "miRBase: Tools for microRNA genomics," *Nucleic Acids Res.*, 2008, doi: 10.1093/nar/gkm952.

[24] L. Fu, B. Niu, Z. Zhu, S. Wu, and W. Li, "CD-HIT: Accelerated for clustering the next-generation

sequencing data," *Bioinformatics*, 2012, doi: 10.1093/bioinformatics/bts565.

[25] S. F. Altschul, W. Gish, W. Miller, E. W. Myers, and D. J. Lipman, "Basic local alignment search tool," *J. Mol. Biol.*, 1990, doi: 10.1016/S0022-2836(05)80360-2.
